# Ultrahigh Resolution Structure of the Heat-stable Form-IAq RuBisCO from the Thermophilic Purple Sulfur Bacterium *Thermochromatium tepidum*

**DOI:** 10.1101/2025.02.09.637349

**Authors:** Shenghai Chang, Weiwei Wang, Michael T. Madigan, Long-Jiang Yu, Haichun Gao, Zheng-Yu Wang-Otomo, Xing Zhang, Jing-Hua Chen

## Abstract

Ribulose 1,5-bisphosphate carboxylase/oxygenase (RuBisCO) catalyzes the initial carbon fixation reaction in the Calvin-Benson-Bassham cycle in plants and most microbial autotrophs. Among the many forms of RuBisCOs, form-I—a protein complex containing 8 large and 8 small subunits—is the most common, representing over 90% of all known RuBisCOs. Although many form-I RuBisCO structures have been determined, no structure has been reported for a form-IAq RuBisCO. Here, and at ultrahigh (1.55 Å) resolution, we detail the structure of the heat-stable form-IAq RuBisCO from the thermophilic and anaerobic purple bacterium *Thermochromatium* (*Tch.*) *tepidum*. The overall structure of the *Tch. tepidum* form-IAq RuBisCO resembles both a form-IAc RuBisCO from a chemolithotrophic sulfur bacterium and a synthetic form-I RuBisCO reconstructed from ancestral sequences. However, despite significant structural and sequence similarities with other form-I RuBisCOs, the *Tch. tepidum* form-IAq RuBisCO shows significantly greater interactions between adjacent small subunits through their extended N-terminal domains that contain a characteristic six-residue insertion unique to form-IAq RuBisCOs. Structural differences of *Tch. tepidum* RuBisCO from its mesophilic relative *Allochromatium vinosum* suggests the mechanisms of its enhanced thermostability. In addition to small subunit interactions, key substitutions on the hydrophilic surface of the small subunits likely contribute to its thermostability as well. It has been proposed that form-I RuBisCOs first appeared in anaerobic thermophiles. Our structure of the *Tch. tepidum* RuBisCO represents the first such structure of a form-IAq enzyme, providing fresh clues for unraveling the evolutionary history of RuBisCO and new details for how this key enzyme remains active at elevated temperatures.

## Introduction

RuBisCO is the key enzyme of carbon fixation through the Calvin-Benson-Bassham cycle in plants and many microbial autotrophs and converts CO_2_ to organic form, contributing over 90% of global carbon fixation^1,2^. RuBisCOs are distributed in all domains of life and are classified into four groups—forms Ⅰ, Ⅱ, Ⅲ and IV—based on their amino acid sequences and structural properties^3,4^. Form-I RuBisCOs are most common and are present in autotrophic *Proteobacteria*, cyanobacteria, algae, and higher plants. These enzymes have a hexadecameric (L_8_S_8_) structure consisting of eight 55-kDa subunits and eight 15-kDa subunits, referred to as large (RbcL) and small (RbcS) subunits, respectively^5–9^. Compared with form-I enzymes, forms II–IV RuBisCOs lack RbcS subunits and exist in various large-subunit-only (L_2_)_n_ complexes that exhibit distinct catalytic activities and activation modes^10^. Form-IV, also known as “RuBisCO-like proteins”, are most divergent and were discovered in the phototrophic green sulfur bacterium *Chlorobaculum tepidum* and endospore-forming bacterium *Bacillus subtilis* where they function in methionine biochemistry rather than CO_2_ fixation^11,12^.

As an important enzyme, the evolution of RuBisCO has been debated, and when and how a RbcS subunit was first added to an (L_2_)_n_-type RuBisCO to yield a primitive form-I heterocomplex has been the subject of much speculation^13–17^. Recent analyses of metagenome-assembled genomes (MAGs) have identified a new clade of form-I RuBisCOs termed “form-I Anaero” that encodes RbcS subunits and is hypothesized to be an ancestral form of the enzyme^14^. The RuBisCO-encoding MAGs were obtained from hot spring environments and belong to organisms related to anaerobic, thermophilic *Bacteria*^14^; notably, these RuBisCOs branch close to the last common ancestor of all known form-I RuBisCOs^14^.

Form-I RuBisCOs are encoded by the genes *cbbL* (large subunit) and *cbbS* (small subunit) and can be further categorized into four sub-forms: IA (found in *Proteobacteria* and *Cyanobacteria*), IB (*Cyanobacteria* and *Prochlorales*), IC (*Proteobacteria* and *Chloroflexi*), and ID (*Proteobacteria* and *Eukaryotes*) based on sequence homology of the proteins^18^. Biochemical studies of purple bacteria—model phototrophs for the study of photosynthesis—have shown that any of three forms of RuBisCO (I, II, IV) or variants (Aq, Ac, or C) of form-I can be present in these organisms with the genomes of many species encoding two or three forms or form-I variants (Supplementary Table 1). Although many of these enzymes have been isolated and biochemically characterized^19^, only a handful of RuBisCO structures from these phototrophs are available and none from purple sulfur bacteria, the preeminent autotrophs of the purple bacterial group (Supplementary Table 1). For example, the purple bacterium *Rhodospirillum* (*Rsp.*) *rubrum* is well known for its form-II homodimeric (L_2_) RuBisCO whose unbound and RuBP-bound structures were determined at resolutions of 1.7 Å and 2.6 Å, respectively^20,21^. The structure of a form-II RuBisCO bound with an activated transition-state analog was reported at 1.85–2.38 Å resolution for the purple nonsulfur bacterium *Rhodopseudomonas* (*Rps.*) *palustris*; this enzyme forms a hexamer composed of three pairs of RbcL homodimers (L_2_)_3_^22^. However, despite being the predominant form of RuBisCO in purple bacteria, the only structure of a form-I RuBisCO (form-IC) from these organisms was reported from *Cereibacter* (*C*., formerly *Rhodobacter*) *sphaeroides* at 3.4 Å resolution^23,24^.

To fill a major gap in our structural understanding of form-I RuBisCOs, here we present the cryo-EM structure of a form-IAq enzyme at ultrahigh resolution (1.55 Å). The enzyme was isolated from the anaerobic and thermophilic purple sulfur bacterium *Thermochromatium* (*Tch.*) *tepidum* (γ-*Proteobacteria*), a phototrophic microbe that grows autotrophically in sulfidic hot springs in Yellowstone National Park, USA^25^. To our knowledge, the structure of the *Tch. tepidum* enzyme is the first form-IAq RuBisCO to be solved at atomic resolution and thus provides the missing piece necessary for completing the evolutionary picture of form-I RuBisCOs. Moreover, because the *Tch. tepidum* RuBisCO is of a “plant-type” and is heat stable^26,27^, its structure may reveal clues to its stability that could someday benefit plant agriculture. As a model purple sulfur bacterium, many *Tch. tepidum* proteins have shown high thermal and structural stability and have yielded crystal structures with higher resolutions than their counterparts from mesophilic species. These include the *Tch. tepidum* reaction center complex (RC, 2.2 Å)^28^, light-harvesting–reaction center complex (LH1–RC, 1.9 Å)^29^, various *c*-type cytochromes^30,31^, and high-potential iron-sulfur protein (HiPIP, 0.48 Å)^32^ [as of late 2024, the latter holds the record for the highest resolution structure among proteins in the Protein Data Bank].

Structural analyses of the *Tch. tepidum* RuBisCO nicely complement these milestones by providing a detailed view of a protein that does not participate in this phototroph’s photosynthetic light reactions but is nevertheless key to its photosynthetic success.

## Results

### Overall structure of the *Tch. tepidum* form-IAq RuBisCO

The *Tch. tepidum* genome contains two sets of form-I RuBisCO genes, one set encoding form-IAq and one set form-IAc^33^ (Supplementary Table 1). From our results here only form-IAq is expressed in cells grown at the optimal temperature (48 °C), and biochemical characterization of this enzyme was accomplished more than three decades ago^26,27^. In the present study, *Tch. tepidum* RuBisCO was highly purified by ammonium sulfate fractional precipitation. SDS-PAGE and gel filtration analyses demonstrated that *Tch. tepidum* RuBisCO was composed of two kinds of subunits, a 52.6-kDa large subunit and a 13.8-kDa small subunit, with a total molecular weight of about 530 kDa (Supplementary Fig. 1). The cryo-EM structure of the *Tch. tepidum* RuBisCO was determined at a global resolution of 1.55 Å (Supplementary Table 2 and Supplementary Fig. 2). Based on the high-resolution density map, side chains of most residues were sufficiently resolved to identify them and confirm that the purified RuBisCO was indeed form-IAq; the final *Tch. tepidum* RuBisCO structure verified this conclusion.

The overall structure of the *Tch. tepidum* form-IAq RuBisCO consists of eight RbcL subunits and eight RbcS subunits, yielding a hexadecamer with a 422-symmetry like other form-I RuBisCOs (Fig. 1a). The four large-subunit dimers assemble into a barreled core forming a solvent accessible channel that runs through the interior (Fig. 1a, Supplementary Fig. 3). Each RbcL subunit contained an N-terminal domain (residues 1–140) and a C-terminal domain (residues 141–472), and two RbcL subunits combine head (N-terminal) to tail (C-terminal) to form a functional dimer (Fig. 1c). The catalytic site of *Tch. tepidum* RuBisCO is formed by residues Lys^167^, Lys^169^, Lys^193^, Asp^195^ and Glu^196^ from the C-terminal domain of one RbcL subunit and three residues (Glu^52^ and Asn^115^) from the N-terminal domain of the adjacent RbcL subunit (Fig. 1c); these were expected because residues near the active site of form-I RuBisCOs are highly conserved^3^ (Fig. 2a). Due to insufficient densities, the C-terminal domain from Phe^459^ to Lys^472^ in the *Tch. tepidum* RbcL subunit was invisible in the density map. Moreover, no densities corresponding to the Mg^2+^ or to substrates were observed within the catalytic site, and thus the structure described here is of the protein’s apo state (Fig. 1c). A total of 1,561 water molecules were modelled into the *Tch. tepidum* RuBisCO structure (Supplementary Fig. 4). Water distribution was not uniform, with more molecules located on the interface between the RbcL dimers and eleven molecules positioned around the catalytic site (Supplementary Fig. 4c). The four RbcS subunits in the *Tch. tepidum* form-IAq RuBisCO were positioned in the usual fashion for form-I enzymes but exhibited extensive intersubunit interactions through their extended N-terminal domains (Supplementary Fig. 3c); the latter structural feature of form-IAq RuBisCOs is distinct from that in other form-I RuBisCOs. In addition to RbcS intersubunit interactions, interactions between RbcL and RbcS subunits were also apparent, and all interactions are discussed in detail now.

**Figure 1.**
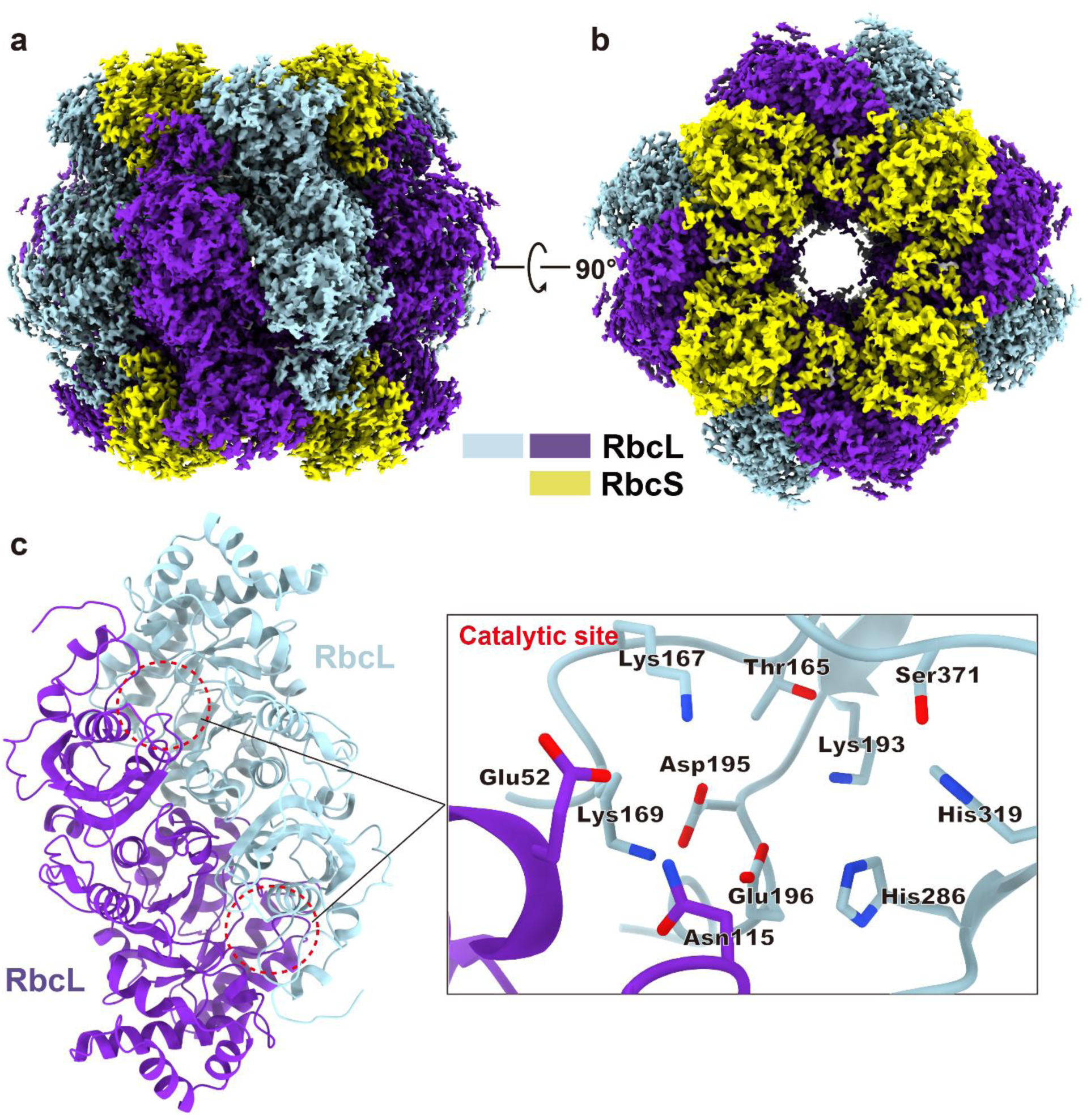
Cryo-EM structure of *Tch. tepidum* form-IAq RuBisCO. **a, b.** The electrostatic potential density map of *Tch. tepidum* form-IAq RuBisCO is shown in side (**a**), and top (**b**) views. **c.** Structure of an RbcL dimer of the *Tch. tepidum* form-IAq RuBisCO shown in cartoon mode. The active sites are marked with red circles and are enlarged in the right panel.

**Figure 2.**
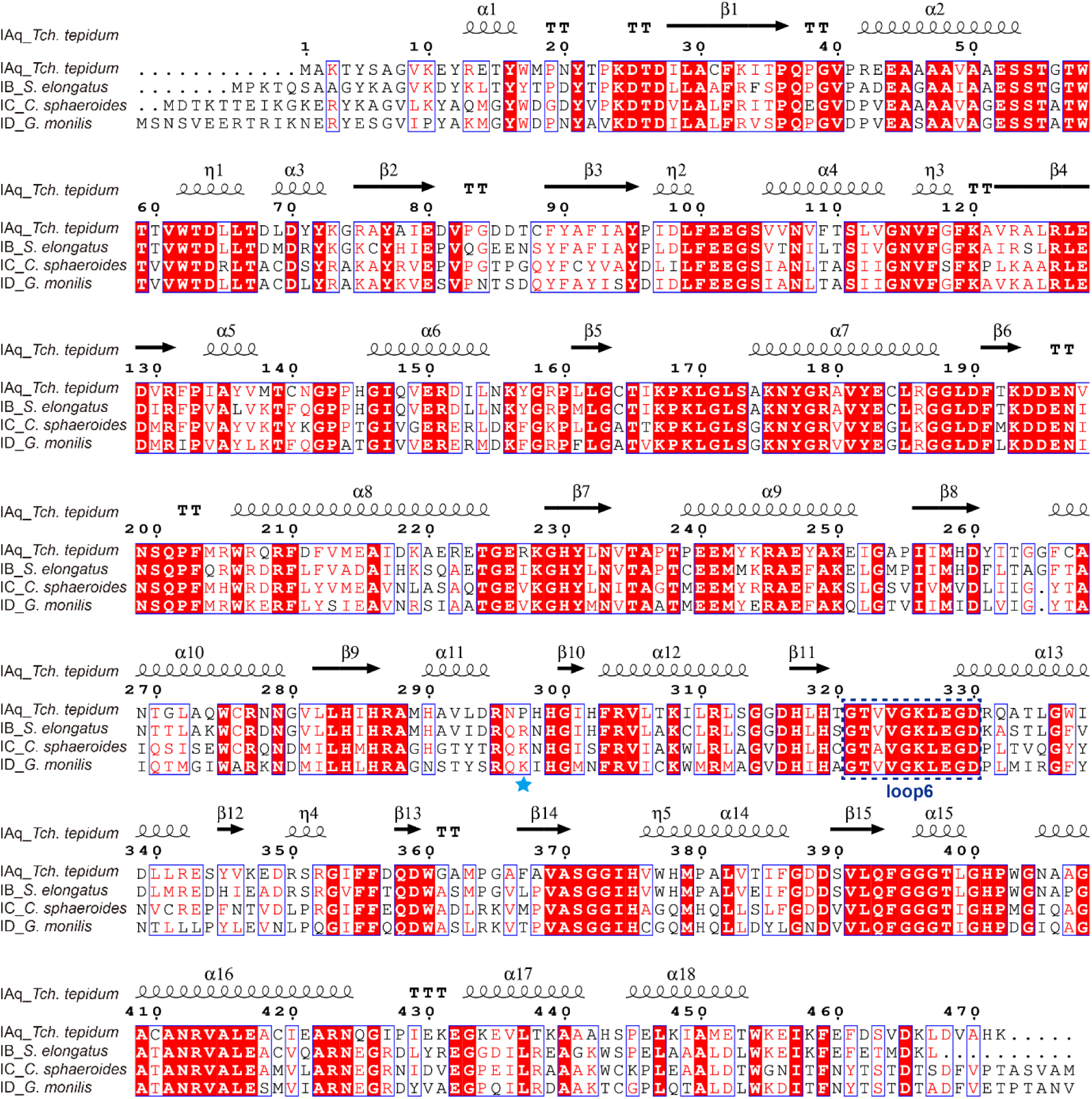
Comparisons of amino acid sequences and structures of the RbcL subunits between *Tch. tepidum* form-IAq RuBisCO and the form-I RuBisCOs of *Synechococcus elongatus*, *Cereibacter sphaeroides* and *Griffithsia monilis*. **a.** Sequence alignments. The residue Pro^297^ and the conserved loop 6 are marked with a cyan pentagram and dashed box, respectively. Residue numbering for sequences and structural annotations (α, α-helix; β, β-strand; η, 3^10^-helix; TT, tight β-turns) are relative to *Tch. tepidum* Rubisco. Alignments were performed using ClustalW^54^, followed by manual curation of output files. Graphics were generated with ESPript^55^. **b.** Structural comparisons. Loop 6 is shown in red. The C-terminal loops are highlighted in yellow.

### Intersubunit interactions in *Tch. tepidum* RuBisCO

The RbcL subunit of *Tch. tepidum* RuBisCO interacts extensively with one RbcS subunit mainly through four regions: Gln^148^–Arg^159^, Arg^186^–Gly^188^, Glu^223^–Arg^227^ and Trp^403^–Gln^425^. The RbcL subunit also interacts with a second RbcS subunit mainly through Gly^171^–Tyr^182^, and several water molecules mediate interactions on the interface between large and small subunits (Fig. 3 and Supplementary Fig. 5). The RbcS subunit of *Tch. tepidum* RuBisCO interacts with one RbcL subunit through its N-terminal domain (Met^3^–Tyr^39^) and C-terminal (Ala^103^–Glu^109^) domain and with a second RbcL subunit in the RbcL dimer through its mid-region (Glu^55^–Lys^66^) (Supplementary Fig. 5).

**Figure 3.**
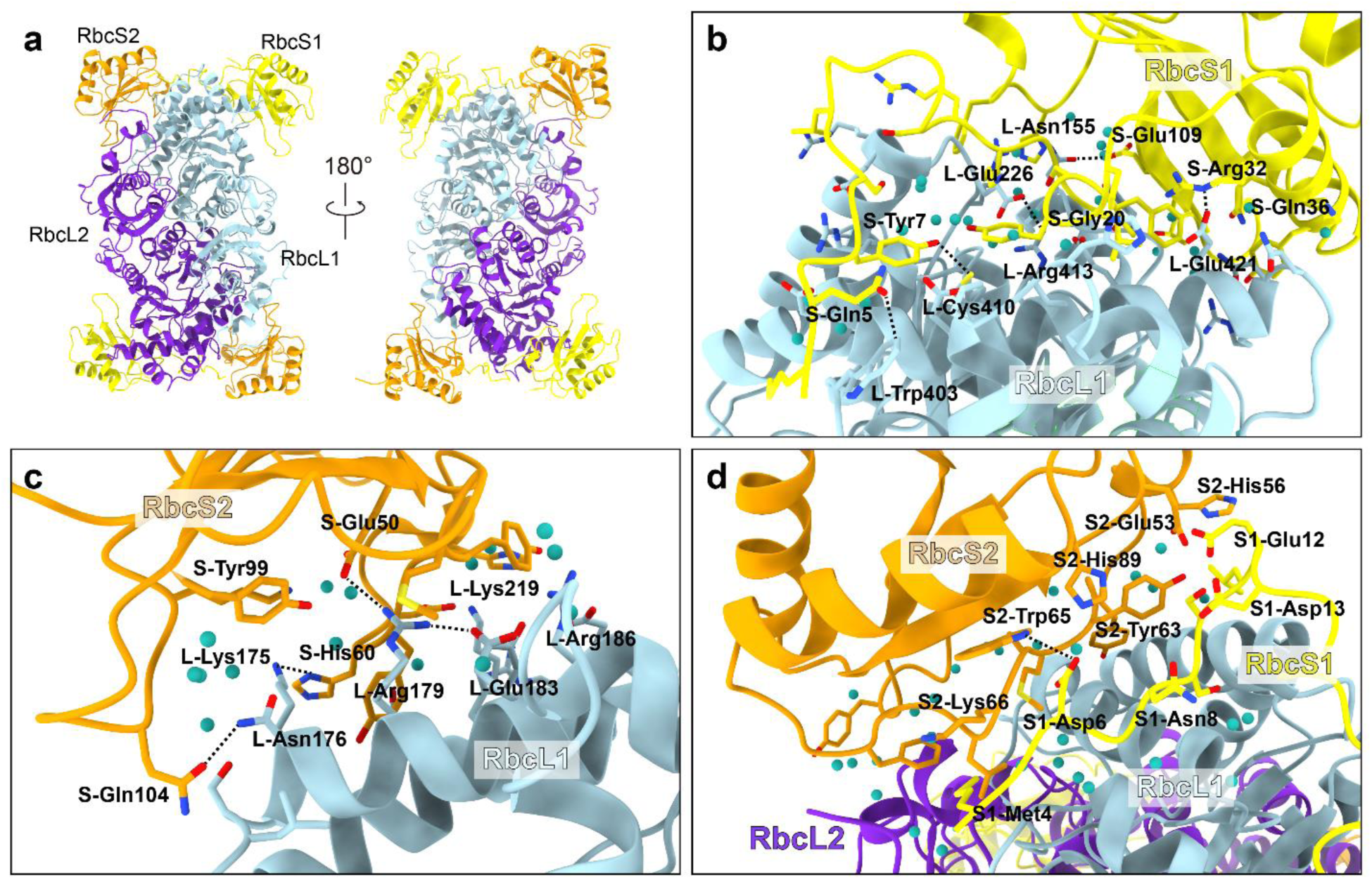
Inter-subunit interactions in the *Tch. tepidum* form-IAq RuBisCO. **a. A** structural unit composed of an RbcL dimer and four associated RbcS subunits. **b.** Close contacts (<3.5 Å, hereafter) between RbsL-1 and RbcS-1. **c.** Close contacts between RbsL-1 and RbcS-2. **d.** Close contacts between RbsS-1 and RbcS-2.

Form-IAq RuBisCOs are characterized by an extended N-terminal domain in the RbcS subunit (Fig. 4 and Supplementary Fig. 5) that contains a six-residue insertion^4^. Each RbcS subunit of the *Tch. tepidum* RuBisCO connects two RbcL dimers and further interacts with an adjacent RbcS subunit through this N-terminal insert (Ser^10^–Asn^15^). This forms an extended loop (Fig. 3d and Supplementary Fig. 5) that helps stabilize the entire RuBisCO molecule. In addition, the N-terminal domain (Met^3^–Glu^12^) of an RbcS subunit interacts with the mid-region (His^51^–Leu^67^) of an adjacent RbcS subunit. Among these, Glu^12^ forms hydrogen bonds with both Glu^55^ and His^56^ in the adjacent RbcS subunit (Fig. 3d). Notably, this Glu^12^ is located in the N-terminal extended loop formed by the unique six-residue insert present in form-IAq RuBisCOs (Fig. 4 and Supplementary Fig. 5).

**Figure 4.**
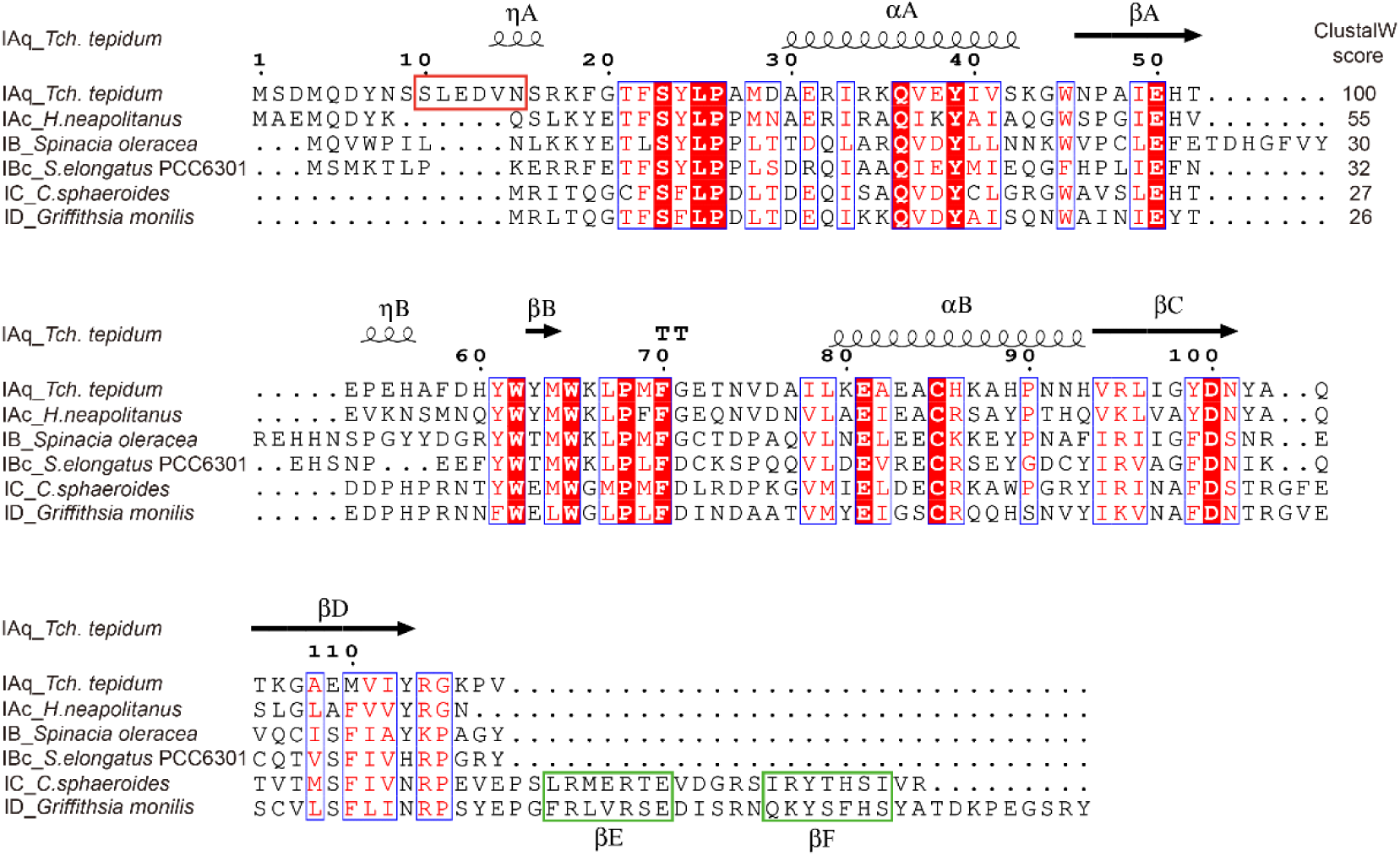
Sequence alignments of the RbcS subunits between *Tch. tepidum* form-IAq RuBisCO and other form-I RuBisCOs. Sequence similarities calculated by ClustalW are indicated on the right side of the top rows. The six-residue insertion in the N-terminal domain of form-IAq RuBisCO is boxed in red, and two additional β-strands in the form-IC and ID RuBisCOs are boxed in green. Sequence and structural annotations are same as in Fig. 2.

### Structural comparison of form-I RuBisCOs between *Tch. tepidum* and other species

Among form-1 RuBisCOs with known structures, the *Tch. tepidum* form-IAq RuBisCO showed the highest structure and amino acid sequence similarities with form-IAc RuBisCO (cloned and expressed in *E. coli*) from the chemolithotrophic and autotrophic bacterium *Halothiobacillus* (*H*.) *neapolitanus* (γ-*Proteobacteria*), a bacterium that grows only on an inorganic sulfur compound (thiosulfate) as the electron donor and CO_2_ as carbon source. A search at the Dali server revealed that the structure of the *Tch. tepidum* form-IAq RuBisCO had the highest Z-scores with the RbcL (PDB: 7SMK) and RbcS (PDB: 6UEW) subunits of the *H*. *neapolitanus* form-IAc RuBisCO (Z-scores of 63 and 20, respectively), corresponding to root mean square deviations (RMSDs) of 0.5 Å and 0.6 Å, respectively^34–36^. These high structural similarities reflect high similarities of amino acid sequences as indicated by ClustalW values of 88 and 55 for RbcL and RbcS subunits, respectively, for *Tch. tepidum* and *H*. *neapolitanus* RuBisCOs (Supplementary Fig. 6).

It is noteworthy that in terms of structural similarity to the *Tch. tepidum* form-IAq RuBisCO indicated by the Dali server, the top group also included a form-I RuBisCO from a sequence reconstruction of the last common RuBisCO ancestor (PDB: 7QSW)^14^. This sequence showed high Z-scores of 58 (RMSD: 0.8 Å) and 18 (RMSD: 1.0 Å) for RbcL and RbcS subunits, respectively, implying a close relationship between the ancestral sequence and *Tch. tepidum* RuBisCO. Sequences of the synthetic constructs were designed based on phylogenetic analysis of the form-I Anaero RuBisCO for the ancestral RbcL (AncLS) and RbcS (AncSSU) subunits^14^. Both AncLS and AncSSU showed high sequence similarities to those of *Tch. tepidum* RbcL and RbcS subunits (Supplementary Fig. 6). The structure of the *Tch. tepidum* RbcL subunit is also similar to that of the RbcL subunits of form-IB RuBisCO from *Synechococcus* (*S.*) *elongatus* (PDB: 1RSC, Z-score: 58), form-IC from *C. sphaeroides* (PDB: 5NV3, Z-score: 51) and form-ID from the red alga *Griffithsia* (*G.*) *monilis* (*Eukarya*) (PDB: 8BDB, Z-score: 55) with Cα RMSDs from 0.9 to 1.2 Å (Fig. 2b). This is consistent with the high sequence identities between the RbcL subunits of *Tch. tepidum*, *S. elongatus*, *C. sphaeroides* and *G. monilis* (Fig. 2a).

Compared with ligand-bound RuBisCOs, a large conformational change was observed for loop 6 in the *Tch. tepidum* apo-RuBisCO (Fig. 2b, Supplementary Fig. 7a and 7b) and slightly different conformations were found for the sidechains of the residues around the catalytic site (Supplementary Fig. 7c and 7d). The flexible loop 6 is composed of ten conserved residues (G^321^TVVGKLEGD^330^) (Supplementary Fig. 6) located close to the catalytic site, and movement of loop 6 is an important mechanism for catalysis and specificity of the enzyme through transitions between open (unliganded or loosely liganded) and closed (tightly liganded) states^37,38^. In this regard, the structure of *Tch. tepidum* form-IAq RuBisCO presented here corresponds to the open state.

### Comparison of the *Tch. tepidum* and *Alc. vinosum* form-IAq RuBisCOs

Compared with RuBisCO from the mesophilic purple bacterium *Alc. vinosum*, the enzyme from *Tch. tepidum* is significantly more thermal stable and shows a higher optimal temperature for activity^26,27^. Assuming because of their relatively high sequence similarities (Fig. 5) that *Alc. vinosum* RuBisCO folds to essentially the same structure as *Tch. tepidum* RuBisCO, sequence comparisons of the two species’ enzymes revealed a much higher occurrence of amino acid substitutions in the *Tch. tepidum* RbcS subunits than in their RbcL subunits (9.3% vs. 3.4%, respectively). This implies that substitutions in the RbcS subunit may contribute more than substitutions in the RbcL subunit to the difference in thermostabilities observed. In the *Tch. tepidum* RuBisCO structure, all 11 substitutions in the RbcS subunit (except for the invisible Asp^3^) and 12 out of the 16 substitutions (two invisible) in the RbcL subunit are located on the hydrophilic surface of the enzyme exposed to solvents (Fig. 7c). Among the substitutions in the *Tch. tepidum* RbcS subunit, Gly^20^, Tyr^99^ and Thr^105^ have close contacts (<3.5 Å) with the RbcL subunit (RbcS-1 in Supplementary Fig. 5), and His^56^ interacts with both an RbcL subunit and an adjacent RbcS subunit (labeled RbcS-1 and RbcS in Supplementary Fig. 5). These results suggest that these substitutions create subtle structural adjustments in the *Tch. tepidum* RuBisCO that increase favorable (and/or reduce unfavorable) subunit interactions and optimize interactions between the enzyme and solvent necessary to remain catalytically robust at the temperature of this phototroph’s hot spring habitat.

**Figure 5.**
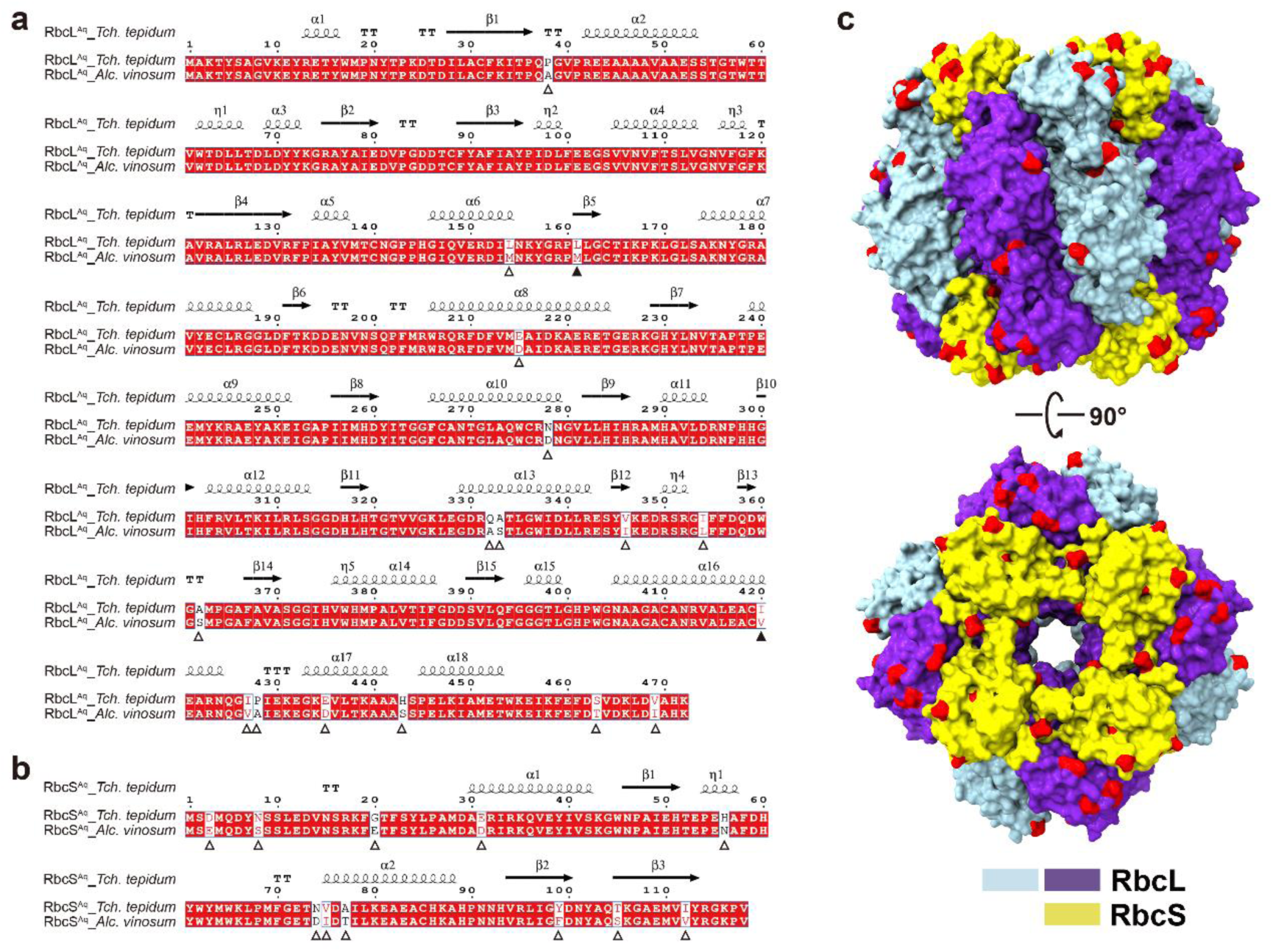
Comparison of the amino acid sequences of the *Tch. tepidum* and *Alc. vinosum* RuBisCO proteins. **a, b.** Sequence alignment of the large subunits (a) and small subunits (b) of the *Tch. tepidum* and *Alc. vinosum* RuBisCO proteins. The substitutions distributed on the hydrophilic surface of the subunits are marked with hollow triangles (Δ). Substitutions buried within the subunits are marked with filled triangles (▴). **c.** The distribution of substitutions on the surfaces of the RuBisCO subunits in side and top views. Substitutions are colored in red.

## Discussion

Here we detail the first structure of a form-IAq RuBisCO. The enzyme, purified from the thermophilic phototrophic bacterium *Tch. tepidum* and detailed at ultrahigh resolution, provides the last missing piece in the structures of known RuBisCOs. The overall structure of *Tch. tepidum* form-IAq RuBisCO most closely resembles the form-IAc RuBisCO from the chemolithotroph *H*. *neapolitanus* and also that of a reconstructed ancestral RuBisCO from a hot spring bacterium. The former is unsurprising because both *Tch. tepidum* and *H*. *neapolitanus* are γ-*Proteobacteria*, whereas the latter may reflect an underlying structural relationship between thermostable RuBisCOs in general and is thus intriguing in terms of both the evolution of form-I RuBisCOs and the structural prerequisites for thermal-stable forms of this enzyme.

The fashioning of RuBisCO from its last common ancestor was accomplished by phylogenetic analysis of a form-I Anaero RuBisCO large (AncLS) subunit reconstructed from a metagenome-assembled genome (MAG). It is hypothesized that this large subunit eventually gained an ancestral small subunit (AncSSU) to yield a primitive form-I (L_8_S_8_) heterocomplex^14^. Extant form-I Anaero RuBisCOs have been detected in anaerobic, thermophilic bacteria of the phylum *Calditrichaeota* that inhabit iron-rich hot springs^39,40^. The amino acid sequence of a form-I Anaero RuBisCO (GenBank: RMG64267.1) encoded from the MAG of a species of *Calditrichaeota* inhabiting a hot spring in Jinata Onsen, Japan revealed relatively high sequence similarities to the *Tch. tepidum* form-IAq RuBisCO for both RbcL and RbcS subunits (Supplementary Fig. 6), signaling a relationship between form-IAq and form-I Anaero RuBisCOs. Because form-I RuBisCOs are thought to have an anaerobic, thermophilic origin^41^, and no structure of a native form-I Anaero RuBisCO exists, the *Tch. tepidum* RuBisCO structure detailed herein is currently the best model of a primitive form-I RuBisCO available.

Compared with other form-I RuBisCOs, form-IA RuBisCOs are unique in possessing an N-terminal extension in their RbcS subunits (Supplementary Fig. 6), and form-IAq RuBisCOs contain a unique six-residue insertion in the RbcS N-terminal region absent from form-IAc RuBisCOs^4^ (Fig. 4). Notably, the structure of the thermostable *Tch. tepidum* form-IAq RuBisCO shows extensive interactions between the adjacent RbcS subunits, a feature not seen to this extent in other form-I RuBisCOs (Fig. 6). For the most part, these interactions occur through residues in the six-residue insertion that form an extended loop on the top exterior face of the protein (Fig. 3d and Fig. 6, Supplementary Fig. 3c). Interactions between this insert and an adjacent RbcS subunit were previously predicted but have now been documented in our work^4^. Sequence comparisons of the *Tch. tepidum* and *Alc. vinosum* RbcS subunits (Fig. 5) bolster the conclusion that substitutions in the *Tch. tepidum* RbcS evolved to exploit interactions between the extended loop and RbcS subunits and thereby stabilize the enzyme to elevated temperatures.

**Figure 6.**
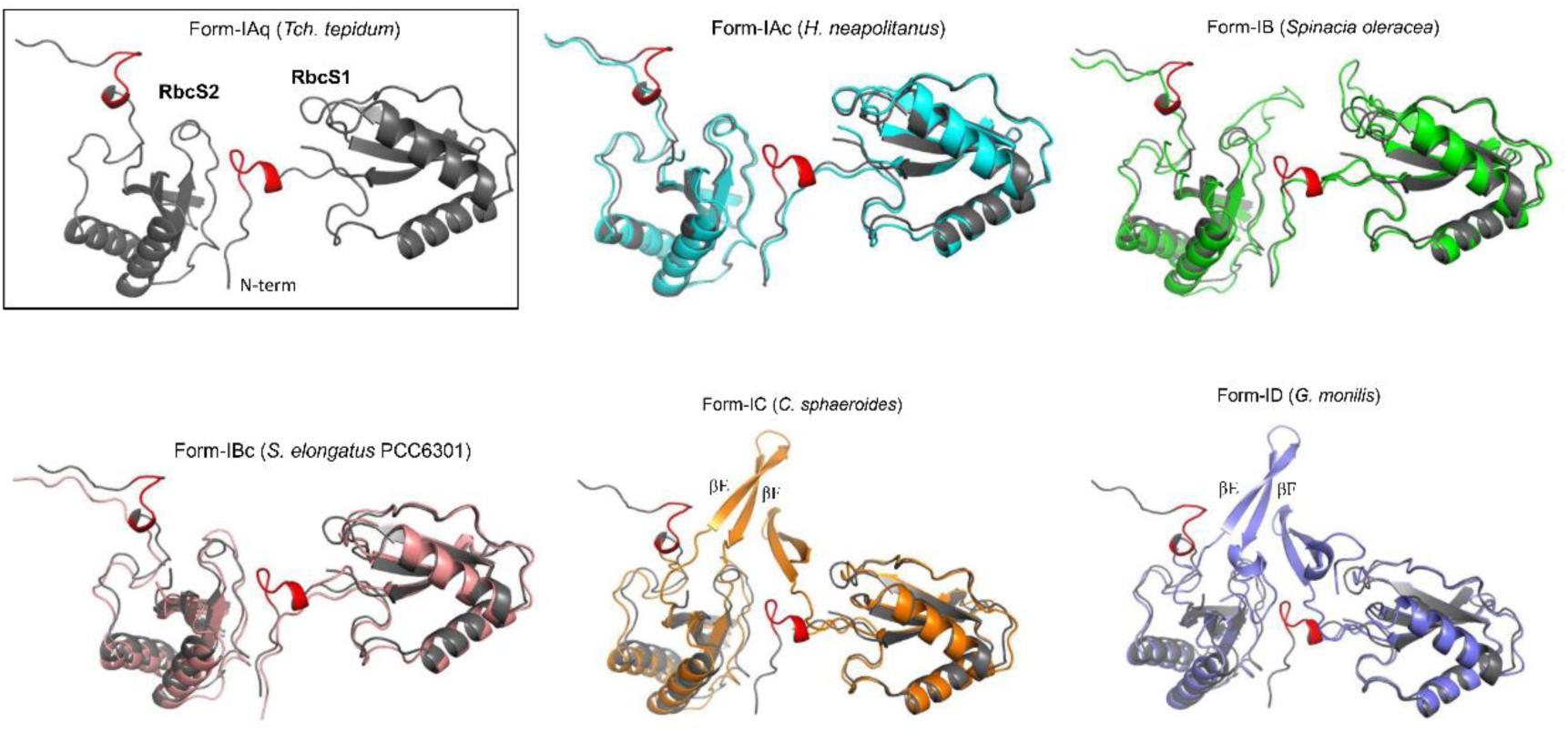
Structural comparisons of the two adjacent RbcS subunits (RbcS1 and RbcS2) between *Tch. tepidum* form-IAq RuBisCO and other representative form-I RuBisCOs. Individual structure of the *Tch. tepidum* RbcS pair is shown in the box. The six-residue insert in the N-terminal domain is shown in red. All other structures are superimposed with that of *Tch. tepidum* (gray). Form-IAc: *H. neapolitanus* (cyan, PDB: 7SMK), form-IB: *Spinacia oleracea* (green, PDB: 1RXO), form-IBc: *Synechococcus elongatus* PCC6301 (light pink, PDB: 1RSC), form-IC: *Cereibacter sphaeroides* (orange, PDB: 5NV3), form-ID: *Griffithsia monilis* (slate blue, PDB: 8BDB). βE and βF indicate two additional β-strands in the form-IC and ID RbcS subunits.

Although likely critical to heat stability, RbcS subunit interactions may not be the sole mechanism responsible for the heat stability of the *Tch. tepidum* RuBisCO. It has been shown that many other *Tch. tepidum* proteins are thermostable with revealed mechanisms. For example, although amino acid identity between high-potential iron-sulfur proteins (HiPIPs) of *Tch. tepidum* and *Alc. vinosum* is about 90%^42^, *Tch. tepidum* HiPIP showed enhanced thermostability attributed to subtle sequence differences that affected the protein’s overall structure^42–44^. Moreover, studies of *Tch. tepidum* cytochrome *c*′ and flavocytochrome *c* revealed several other factors that contributed to the thermostabilities of these soluble proteins^30^ including differences in (i) the number of hydrogen bonds and charged and polar residues, (ii) the number of residues with shorter sidechains (Gly and Ala), and (iii) the distribution of water molecules on both the surface and interior of the proteins. One or more of these factors may also contribute to thermostability of *Tch. tepidum* RuBisCO.

Since the *Tch. tepidum* RuBisCO is a “plant-type” (form-I) enzyme, the RbcS interactions revealed here may provide useful clues for engineering heat stability into plant-type RuBisCOs. Although it is clear that net CO_2_ assimilation in plants is heat sensitive^45^, thus far surprisingly little structural information has been published on heat-stable form-I RuBisCOs, as most studies have focused on the dimeric form-III enzymes from hyperthermophilic *Archaea* that function above 100 °C^46^. However, as Earth’s climate warms, heat-active plant RuBisCOs—especially ones like the *Tch. tepidum* enzyme that functions optimally at mildly elevated temperatures—will eventually be essential to maintain agricultural productivity at the levels required to feed future human and animal populations. Thus, besides filling the last remaining gap in our understanding of RuBisCO structures, details of the *Tch. tepidum* enzyme may have applications for creating heat-tolerant plant RuBisCOs.

## Materials and Methods

### Purification of *Tch. tepidum* RuBisCO

Cells of *Tch. tepidum* were grown anaerobically for 7 days at 49 ℃ with continuous illumination. Ten liters of cells were collected at 4,000 g for 10 min, and the pellet was resuspended in 100 ml Tris buffer (20 mM Tris-HCl, pH 8.0; 1 mM MgCl_2_; 20 ug/ml DNase I). After pre-cooling on ice for 20 min, cells were broken using an ultrasonic crusher (TOMY, UD-211). The lysate was centrifuged at 4 ℃ at 17,000 *g* for 20 min and the pellet discarded. The supernatant was then ultracentrifuged at 150,000 *g* at 4 ℃ for 70 min. The resulting supernatant was filtered through a 0.45 μm membrane filter before undergoing an anion exchange column chromatography (Q-Sepharose High performance (GE)) using an AKTA-FPLC protein purification system. After loading the sample, the column was washed first with Tris buffer containing 160 mM NaCl and then with a gradient elution with the concentration of NaCl increasing from 160 to 300 mM. The sample eluted with 250-270 mM NaCl was collected and concentrated using 30 kDa ultrafiltration tubes (1,000 *g*, 20 min). Subsequently, the sample was precipitated with 50% and 65% ammonium sulfate successively. The samples precipitated with 65% ammonium sulfate were resuspended with Tris buffer (20 mM Bis-Tris-HCl, pH 6.0) and loaded on a second anion exchange column. The column was eluted with a gradient of 150–300 mM NaCl. The first elution peak was collected, concentrated and further isolated by size exclusion chromatography (Superose 6 Increase 10/300 (GE)) with Tris buffer containing 200 mM NaCl. A high-purity RuBisCO sample was eluted with a sharp peak around 15 ml that was collected and concentrated with 100 kDa ultrafiltration tubes (Millipore) to a suitable final concentration for cryo-EM analysis.

### Cryo-EM sample preparation and data collection

To achieve a high-resolution reconstruction, concentrated *Tch. tepidum* RuBisCO proteins of 50 mg/ml were plunge-frozen in liquid ethane using a Vitrobot (FEI) and the vitrified sample was subsequently imaged on a 300 kV FEI Titan Krios cryo-electron microscope equipped with a Falcon4 detector and a Selectris (TFS) image filter. Using EasyGlow (Pelco) was used to glow the grids (Quantifoil Cu1.2/1.3 300 mesh) and make the surface hydrophilic to facilitate an even layer on the grid surface. The glow current intensity was set to 20 mA, and the glow time was 2 min. An aliquot of 3 μL purified *Tch. tepidum* RuBisCO complex sample (50 mg/ml) was applied to a treated grid and then the grids were blotted for 4 s at a humidity of 100% and 20 °C and plunge-frozen in liquid ethane using a Vitrobot (TFS).

Using EPU, cryo-EM images of *Tch. tepidum* RuBisCO were recorded at an FEI Titan Krios electron microscope operated at 300 kV under a nominal magnification of 270,000. The microscope was carefully aligned before data collection, including the coma-free alignment to minimize the effects of beam tilt. The dose rate of the electron beam was set to ∼24 e^-^/Å^2^/s, and movies were recorded on a Falcon4 detector equipped with Selectris (TFS) for 3 s with a total dose of ∼72 e^-^/Å^2^ on the specimen using EPU (TFS).

### Cryo-EM image processing

MotionCor2 was used to correct drift for each image, and using Gctf to measure the contrast transfer function of the corrected images^47,48^. Relion 4.0 and cryoSPARC3.1 were used to complete subsequent data processing^49,50^. Through the autopick function in Relion, 2,137,842 particles were selected from 6,571 images and then cryoSPARC was used for the first 3D classification, resulting in 907,092 particles. An additional round of 3D classification was performed on these particles using Relion, resulting in 765,738 particles. After subjecting these particles to 3D refinement, CTF refinement, and polishing in Relion, and imposing the *D*4 axis of symmetry, a 3D map sharpened at a B-factor of −25 Å² was obtained at a resolution of 1.55 Å according to the gold standard FSC at 0.143. The specific process is shown in Supplementary Fig. 2.

### Modeling and refinement

The initial model of *Tch. tepidum* RuBisCO was constructed by using the RuBisCO structure from a cyanobacterium (PDB code: 6LRR) as a reference and fitting it into the corresponding density map of *Tch. tepidum* RuBisCO using UCSF Chimera^51^. This was followed by manually replacing and editing the model using Coot to obtain the *Tch. tepidum* atomic model^52^. Subsequently, PHENIX was used to optimize the atomic model, and manually adjust it using Coot to obtain the final RuBisCO atomic model^53^. The relevant model building parameters are shown in Supplementary Table 2.

## Data availability

Cryo-EM maps and atomic coordinates of *Tch. tepidum* RuBisCO have been deposited into Electron Microscopy Data Bank (accession code, EMD-60792) and Protein Data Bank (accession code 9IQO), respectively.

## Acknowledgements

We thank the staff from the Center of Cryo-Electron Microscopy (CCEM), Zhejiang University for their technical assistance on cryo-EM data collection, and Dr. Cheng Ma from the Protein facility, School of medicine, Zhejiang University for his assistance on protein purification. This work was supported by a Natural Science Foundation of Zhejiang Province, China (LR22C010001 to J.-H.C.) and National Natural Science Foundation of China (32100202 to J.-H.C). MTM was supported in part by NASA Cooperative Agreement 80NSSC21M0355.

## Author contributions

J.-H.C, X.Z. and Z.-Y.W.-O. initialized the research. J.-H.C., W.W. and L.-J.Y. purified the protein sample. S.C. prepared the cryo-sample, collected the data, build the atomic model. J.-H.C., X.Z., Z.-Y.W.-O., M.T.M., S.C., W.W. and H.G. analyzed the data, interpreted the structures, prepared the figures and wrote the paper. All authors discussed and commented on the results and the manuscript.

## Supplementary Information

**Supplementary Table 1.**
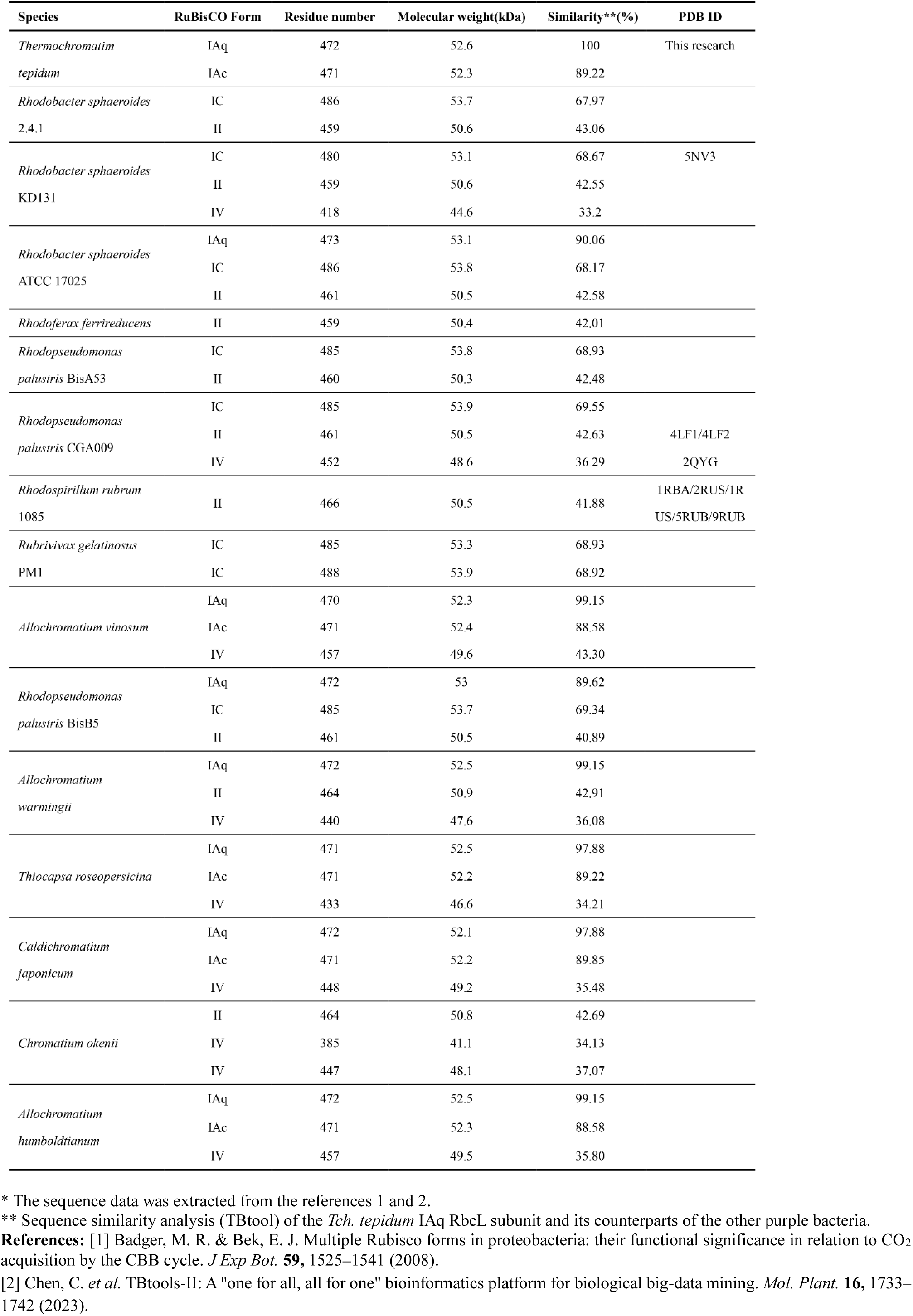
Different forms of RuBisCO proteins identified in purple bacteria*.

**Supplementary Table 2.** Cryo-EM data collection, refinement and validation statistics.

|  | *Tch. tepidum* RuBisCO |
| --- | --- |
| Magnification | 270,000 |
| Voltage (kV) | 300 |
| Electron exposure (e<sup>-</sup>/Å<sup>2</sup>) | 50 |
| Defocus range (um) | -0.3~-1.0 |
| Pixel size (Å) | 0.456 |
| Symmetry imposed | D4 |
| Micrographs | 6,571 |
| Initial particle images(no.) | 2,137,842 |
| Final particle images(no.) | 765,738 |
| Map resolution (Å) (global) | 1.55 |
| FSC threshold | 0.143 |
| Map-sharpening B-factor (Å<sup>2</sup>) | -25 |
| Map resolution range (Å) | 1.4-1.88 |
| **Refinement** |  |
| Initial model used (PDB code) | 6lrr |
| Model resolution | 1.5 |
| FSC threshold | 0.143 |
| Model resolution range (Å) | 1.3-16 |
| **Model composition** |  |
| Non-hydrogen atoms | 37585 |
| Protein residues | 4544 |
| Ligands | 0 |
| **B factors (Å<sup>2</sup>)** |  |
| Protein | 20.73 |
| Ligand | None |
| **R.m.s. deviations (PHENIX)** |  |
| Bond length (Å) | 0.003 |
| Bond angles (°) | 0.541 |
| **Validation** |  |
| MolProbity score | 1.43 |
| Clashscore | 6.78 |
| Poor rotamer (%) | 0 |
| **Ramachandran plot** |  |
| Favored (%) | 97.74 |

**Supplementary Fig. 1.**
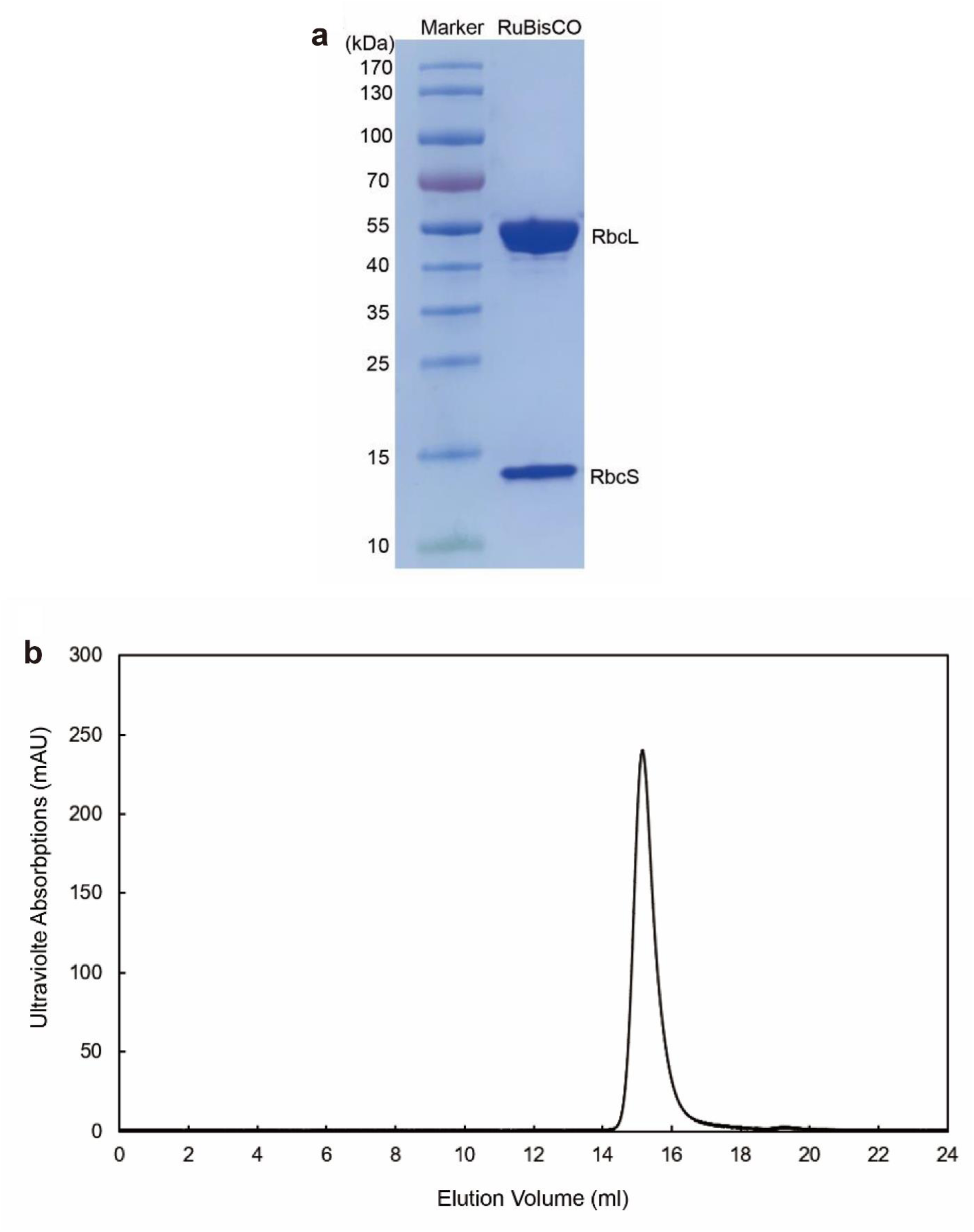
Biochemical analysis of the *Tch. tepidum* form-IAq RuBisCO. **a.** SDS-PAGE analysis of the purified *Tch. tepidum* form-IAq RuBisCO sample with a 4-20% gradient acrylamide gel. **b.** Elution pattern of the purified *Tch. tepidum* form-IAq RuBisCO sample from a gel filtration column Superose 6 Increase (GE Healthcare).

**Supplementary Fig. 2.**
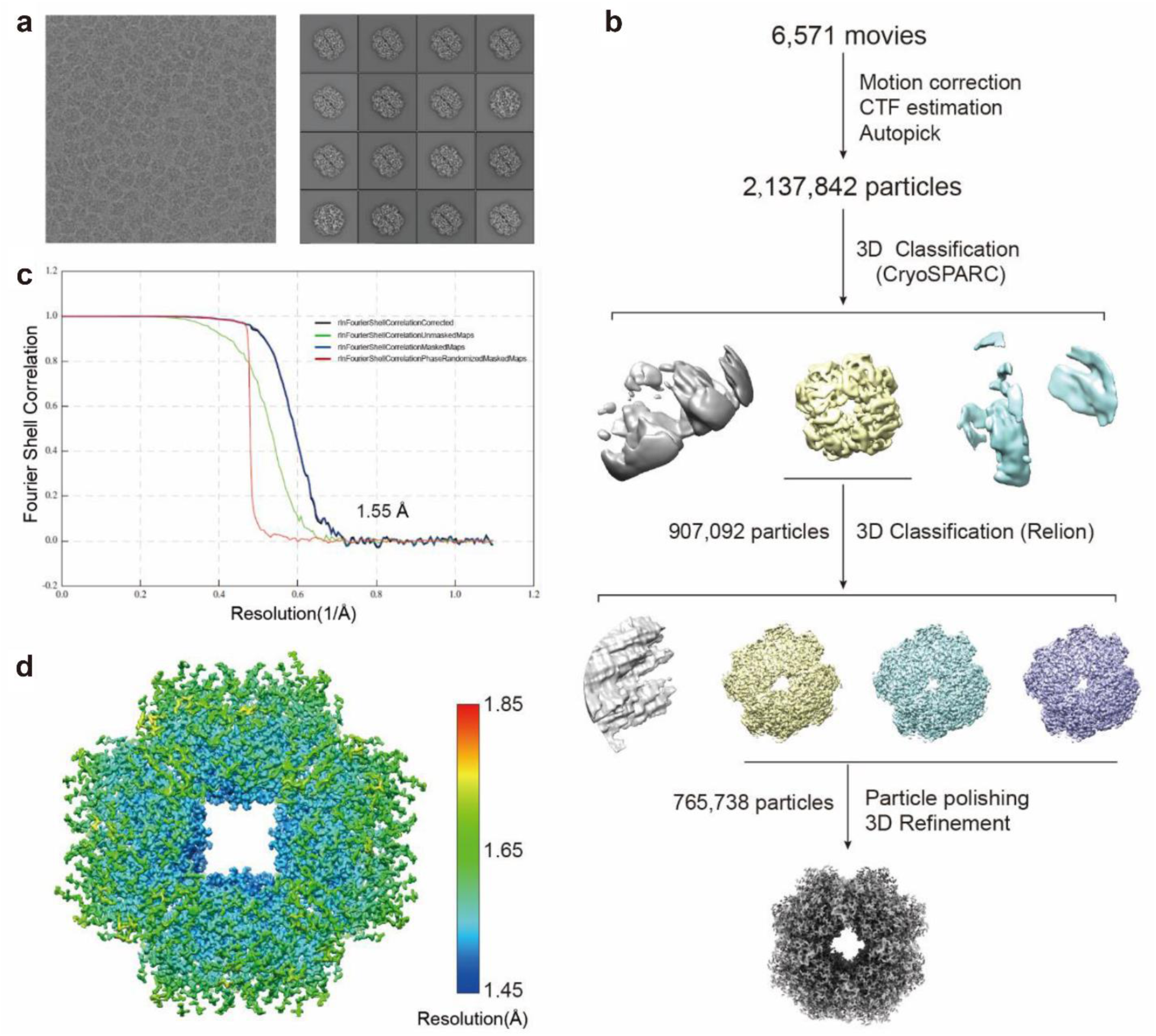
Image-processing flowchart for *Tch. tepidum* form-IAq RuBisCO. **a.** Typical cryo-EM microscope image and the selected class averages from 2D classification of the *Tch. tepidum* form-IAq RuBisCO sample. **b.** Scheme of three-dimensional classification, refinement and local refinement of cryo-EM particle images and the final 3D reconstitution of *Tch. tepidum* form-IAq RuBisCO at 1.55 Å. **c.** Gold-standard FSC curves of the final cryo-EM maps of *Tch. tepidum* form-IAq RuBisCO. **d.** Local resolution distribution of the density map.

**Supplementary Fig. 3.**
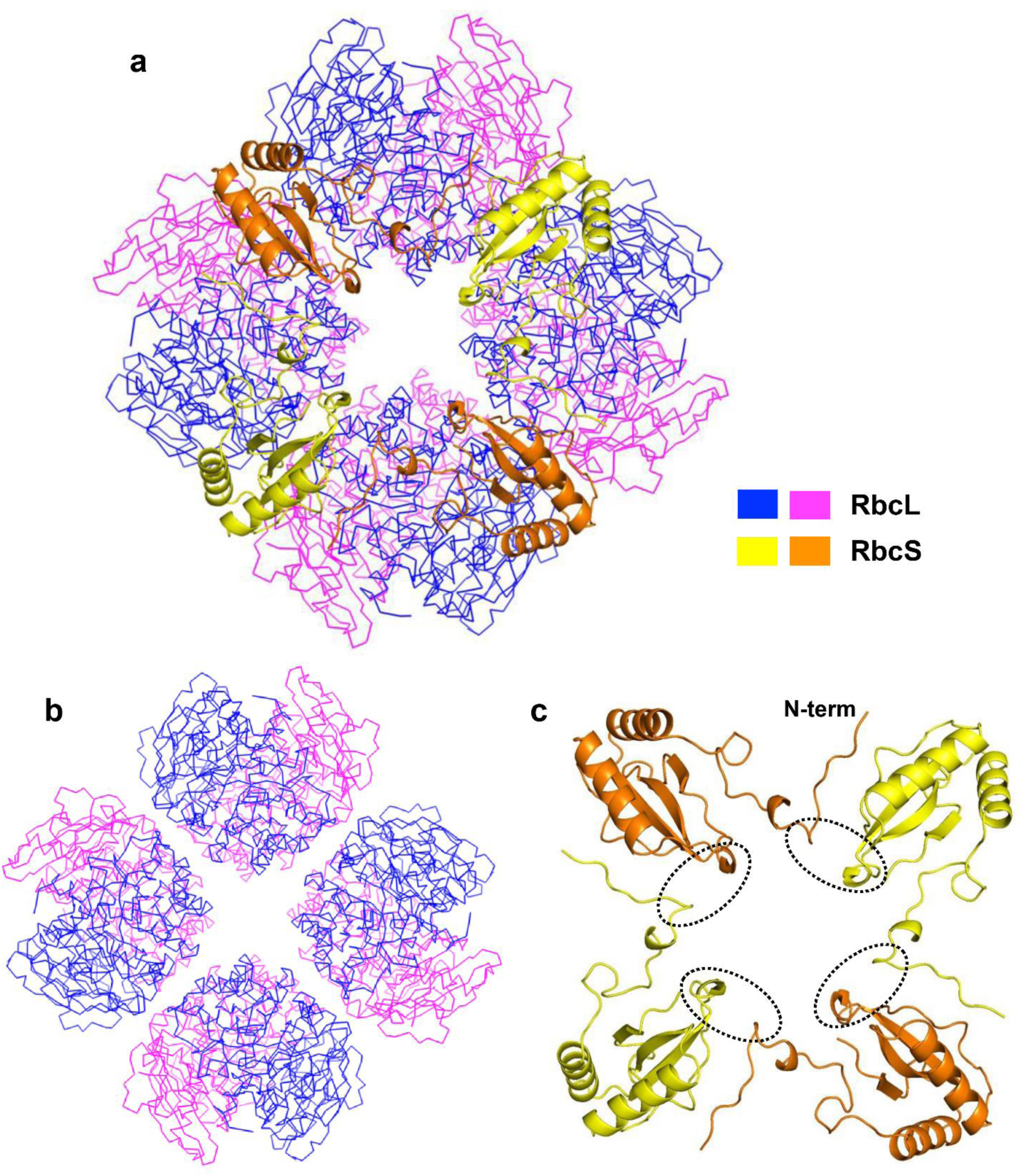
Structure of the Tch. tepidum form-IAq RuBisCO. **a**. Top view of the overall structure. RbcL and RbcS subunits are shown by ribbons and cartoons, respectively. **b.** Top view of the arrangement of the four RbcL dimers. **c.** Arrangement of the four adjacent RbcS subunits. Interactions between the extended N-terminal loop in one subunit and the middle portion in the adjacent subunit are shown by dotted ellipses.

**Supplementary Fig. 4.**
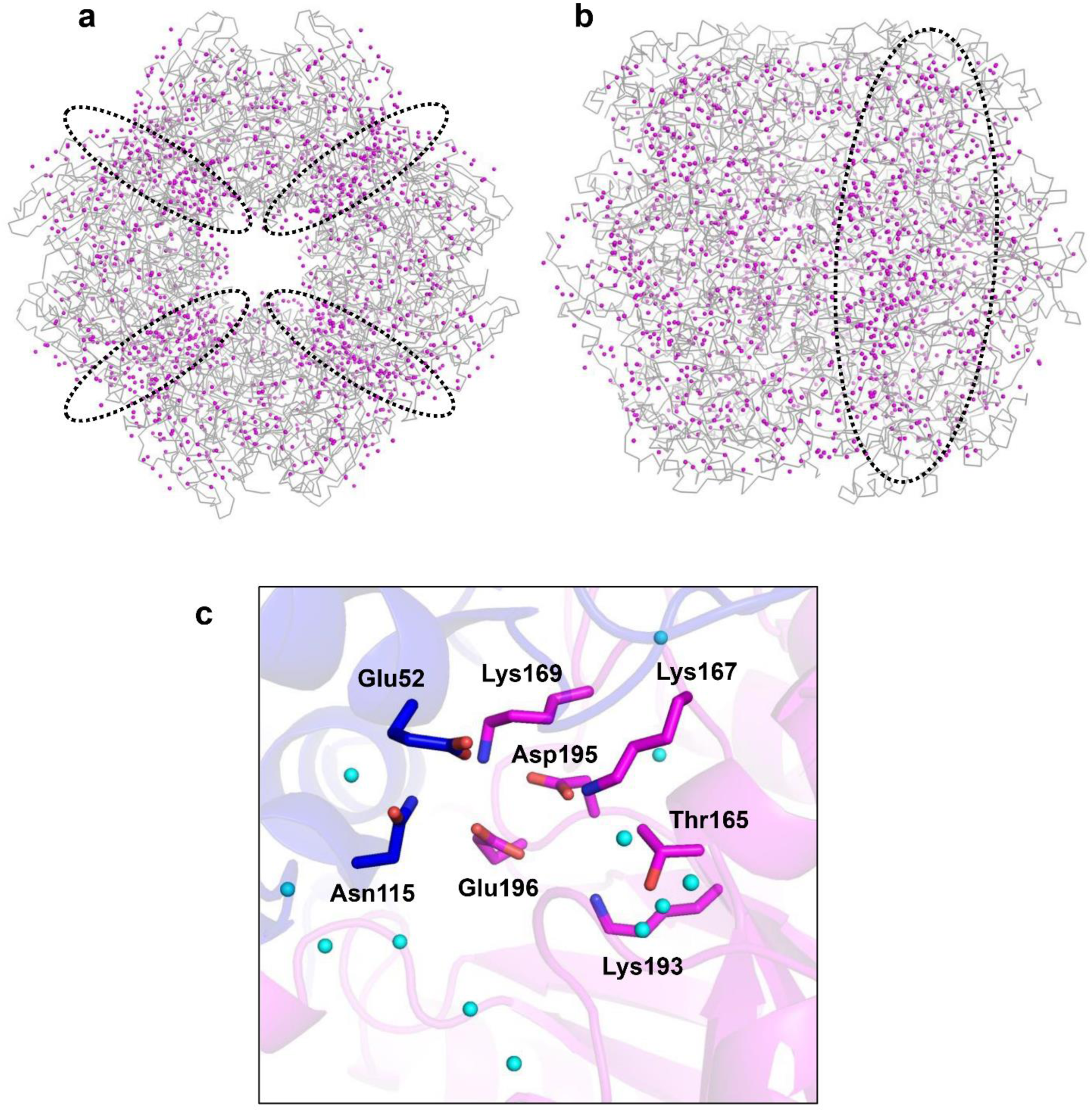
Water molecules in the Tch. tepidum RuBisCO. **a**. Top view of water (magenta dots) distribution with the interfaces between L_2_ dimers indicated by dotted ellipses. **b.** Side view of water (magenta dots) distribution with the interface between L_2_ dimers indicated by a dashed ellipse. **c.** Water molecules (cyan spheres) in the catalytic site.

**Supplementary Fig. 5.**
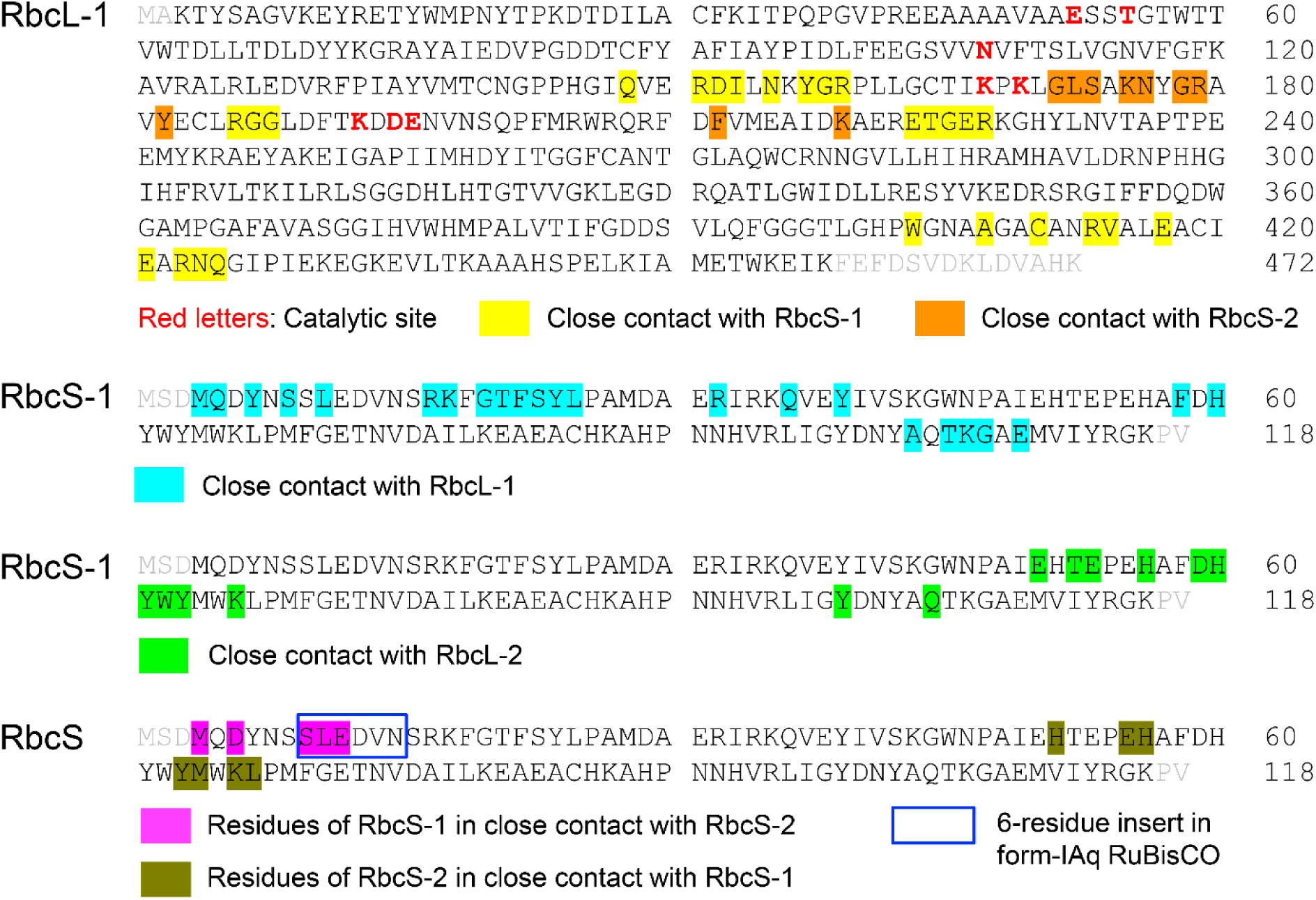
Mapping close contacts (<3.5 Å) between the subunits of *Tch. tepidum* form-IAq RuBisCO. Color-marked letters show the residues interacting between the different subunits as indicated in Fig. 3.

**Supplementary Fig. 6.**
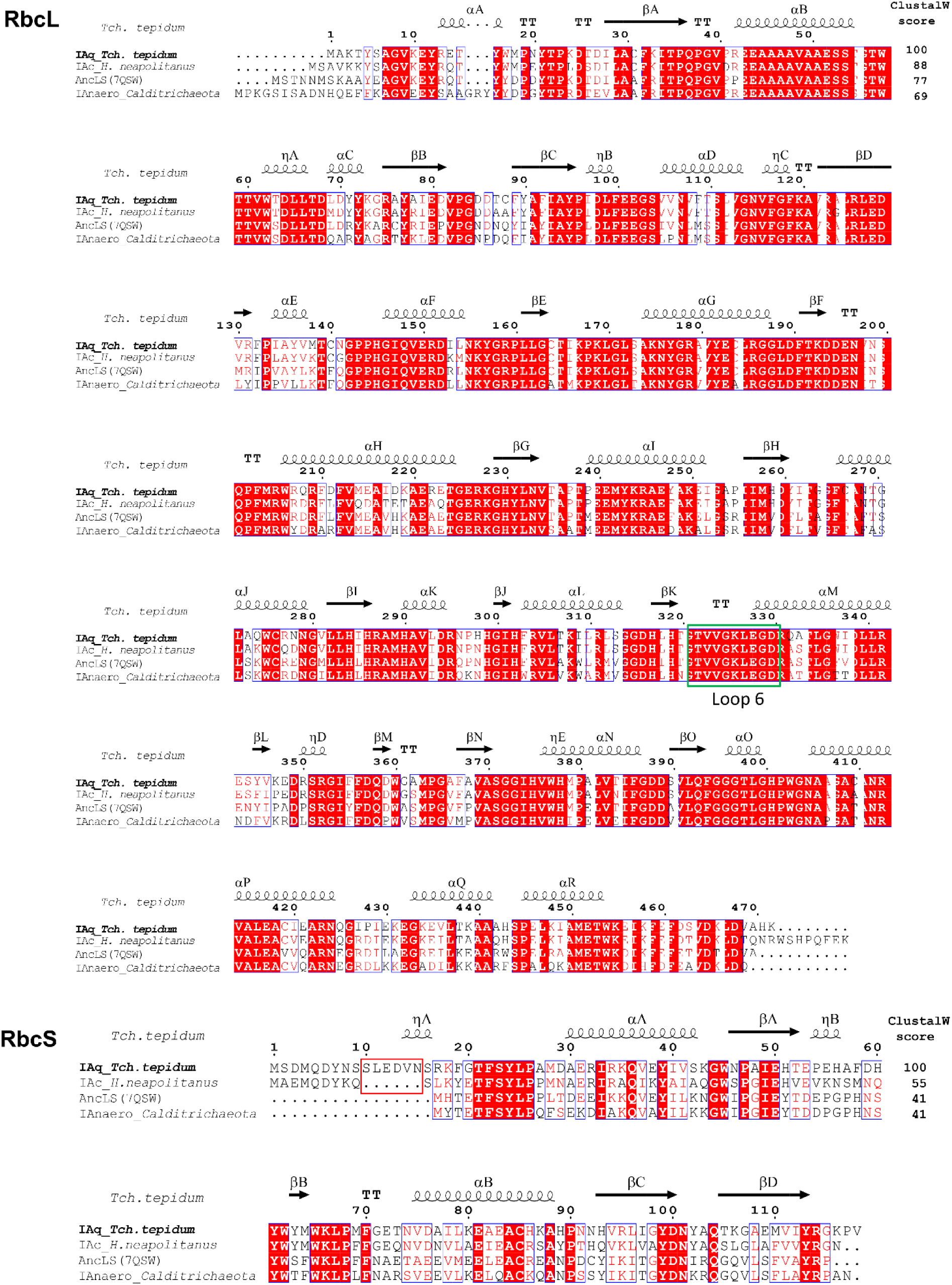
Sequence alignments of the *Tch. tepidum* form-IAq RuBisCO with other form-I RuBisCOs that show high sequence and structural similarities. Sequence similarities calculated by ClustalW are indicated on the right side of the top rows. The Loop-6 in the RbcL subunits is boxed in green and the six-residue insertion in the form-IAq RbcS subunit is boxed in red. Sequence and structural annotations are same as in Fig. 2.

**Supplementary Fig. 7.**
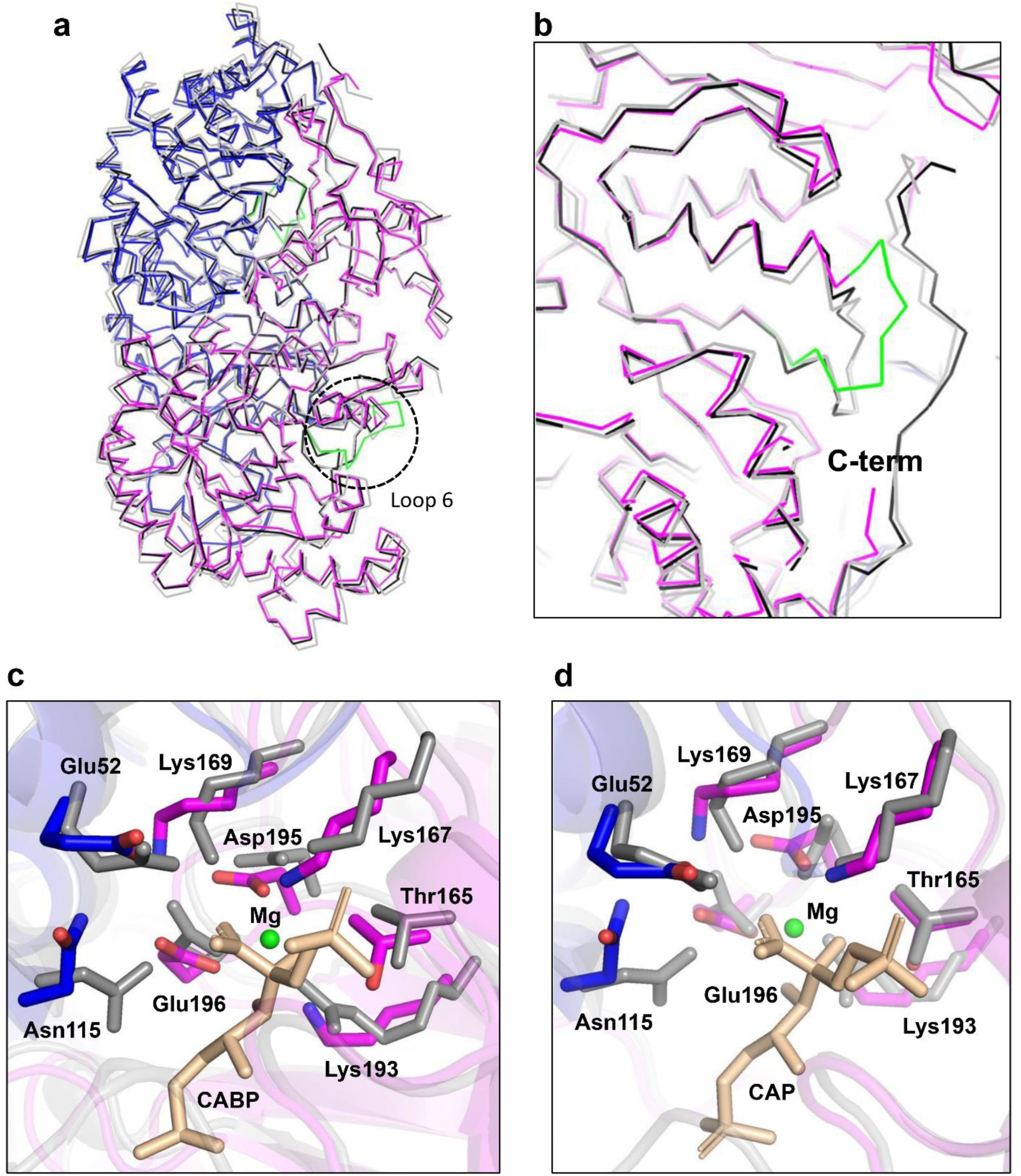
Structural comparisons of the RbcL subunits of *Tch. tepidum* form-IAq RuBisCO with those of other RuBisCOs. **a.** Mainchain superpositions of the RbcL subunit of *Tch. tepidum* (colored) with those of *C. sphaeroides* (gray, PDB: 5NV3) and the reconstructed ancestral RbcL (AncLS) (black, PDB: 7QSW). The loop-6 in *Tch. tepidum* RbcL subunit is colored in green. **b**. Expanded view of the loop-6 (dashed circle) in **a**. **c**. Comparisons of the catalytic sites between *Tch. tepidum* (colored) and *C. sphaeroides* (gray). **d**. Comparisons of the catalytic sites between *Tch. tepidum* (colored) and the reconstructed AncLS (gray).

